# KSHV vIL-6 Enhances Inflammatory Responses by Epigenetic Reprogramming

**DOI:** 10.1101/2023.06.25.546454

**Authors:** Tomoki Inagaki, Kang-Hsin Wang, Ashish Kumar, Chie Izumiya, Hiroki Miura, Somayeh Komaki, Ryan R. Davis, Clifford G. Tepper, Harutaka Katano, Michiko Shimoda, Yoshihiro Izumiya

## Abstract

Kaposi sarcoma-associated herpesvirus (KSHV) inflammatory cytokine syndrome (KICS) is a newly described chronic inflammatory disease condition caused by KSHV infection and is characterized by high KSHV viral load and sustained elevations of serum KSHV-encoded IL-6 (vIL-6) and human IL-6 (hIL-6). KICS has significant immortality and possesses greater risks of having other complications, which include malignancies. Although prolonged inflammatory vIL-6 exposure by persistent KSHV infection is expected to have key roles in subsequent disease development, the biological effects of prolonged vIL-6 exposure remain elusive. Using thiol-Linked Alkylation for the Metabolic Sequencing and Cleavage Under Target & Release Using Nuclease analysis, we studied the effect of prolonged vIL-6 exposure in chromatin landscape and resulting cytokine production. The studies showed that prolonged vIL-6 exposure increased Bromodomain containing 4 (BRD4) and histone H3 lysine 27 acetylation co-occupancies on chromatin, and the recruitment sites were frequently co-localized with poised RNAPII with associated enzymes. Increased BRD4 recruitment on promoters was associated with increased and prolonged NF-KB p65 binding after the lipopolysaccharide stimulation. The p65 binding resulted in quicker and sustained transcription bursts from the promoters; this mechanism increased total amounts of hIL-6 and IL-10 in tissue culture. Pretreatment with the BRD4 inhibitor, OTX015, eliminated the enhanced inflammatory cytokine production. These findings suggest that persistent vIL-6 exposure may establish a chromatin landscape favorable for the reactivation of inflammatory responses in monocytes. This epigenetic memory may explain the greater risk of chronic inflammatory disease development in KSHV-infected individuals.

**Author summary:** Combined and continuous cytokine stimulation triggers transcription reprogramming and is often used for specific tissue development. Continuous vIL-6 exposure occurs in KSHV-infected patients and leads to inflammatory cytokine storm with high mortality. However, possible epigenetic reprogramming by the vIL-6 and its association with pathogenesis remain unclear. Here we demonstrate the establishment of a new chromatin landscape mediated by BRD4 through prolonged vIL-6 exposure which contributes to more robust and rapid transcription and increased cytokines production. Inhibition of BRD4 suppressed this inflammatory response. Our results indicate that targeting the epigenetic effect of viral cytokines may lead to novel therapies for KSHV-induced inflammatory cytokine storms.

## Introduction

Controlled inflammatory response enhances host immunity against external insults, whereas uncontrolled inflammatory responses resulting from excessive cytokine production leads to disruption of such balance. In particular, monocytes and macrophages play major roles in controlling pro-inflammatory cytokines expression and their tightly controlled activation is critical for regulating systemic inflammation in our body. Previous reports have shown that continuous inflammatory stimulation enhances [1, 2] or sometimes impairs [3, 4] responses to subsequent stimulus. In either case, continuous inflammatory stimulation appears to alter the cellular phenotype and affect inflammatory cytokine production.

Kaposi’s sarcoma-associated herpesvirus (KSHV) was first identified in Kaposi’s sarcoma (KS) lesions in 1994 [5]. KSHV is also associated with lymphoproliferative diseases, primary effusion lymphoma (PEL) and multicentric Castleman’s disease (MCD) [6, 7]. Recent studies showed that KSHV infection also causes severe systemic inflammation, which is categorized as Kaposi’s sarcoma-associated herpesvirus inflammatory cytokine syndrome (KICS) [8]. KICS is characterized by increased viral loads and inflammatory cytokines such as human IL-6 (hIL-6), IL-10, and KSHV-encoded interleukin-6 homolog, viral IL-6 (vIL-6). Individuals with KICS are known to have a higher risk of developing KSHV-associated cancers and other malignancies [8].

The vIL-6, encoded by KSHV ORF-K2, is expressed during the lytic replication phase, and single cell transcriptomics studies also suggested that vIL-6 may be expressed during latency in selected cell populations [9]. Similar to hIL-6, vIL-6 is known to enhance cell proliferation, endothelial cell migration, and angiogenesis by upregulating vascular endothelial growth factor (VEGF) [10] and downregulates the caveolin 1 expression [11–13]. Moreover, vIL-6 transduction also leads to cell transformation in the 3T3 cells model and induces tumors in xenograft mice [10]. Increased metastasis is also seen in a murine xenograft model with transgenic vIL-6 mice [14]. The vIL-6 activates downstream signaling pathways by binding to the cellular hIL-6 receptor, gp130. Dimerization of gp130 induced by hIL-6 binding activates Janus tyrosine kinases and phosphorylates the SH2-containing cytoplasmic protein STAT3 (signal transducer and activator of transcription 3). The phosphorylated STAT3 forms a dimer and translocates to the nucleus to activate the downstream genes that play essential roles in inducing the inflammatory response, cell survival, and immune responses [15–17].

Synergistic interactions between another inflammatory-associated transcription factor, nuclear factor-kappa B (NF-κB) and STAT3 are known to induce the hyperactivation of NF-κB followed by the production of various inflammatory cytokines including TNFα and hIL-6. Because hIL-6 expression is regulated by NF-κB pathway activation, simultaneous activation of NF-κB and STAT3 triggers a positive feedback loop of NF-κB activation in the hIL-6/STAT3 axis. This positive feedback loop is called the IL-6 amplifier (IL-6 Amp) and is a key element in inflammatory disease development [18]. In the IL-6 Amp disease model, obesity, injury, and infection trigger chronic inflammation and/or a systemic cytokine storm, which is enhanced by interactions between local non-immune cells and infiltrating immune cells [18]. A prominent example of IL-6 Amp is also seen in Coronavirus disease 2019 (COVID-19), which is triggered by severe acute respiratory syndrome coronavirus 2 (SARS-CoV2) infection [18]. Consistent with the significance of hIL-6 in disease development, a humanized monoclonal antibody against the hIL-6 receptor, tocilizumab, has proven successful for treating Castleman’s disease as well as COVID-19 [19, 20]. In the case of KSHV-associated diseases, the IL-6 Amp may be triggered by KSHV infection and is likely to be associated with vIL-6 expression from KSHV-infected cells. However, how persistent exposure of vIL-6 alters cellular responses and enhances inflammation cytokine production remain elusive.

The bromodomain and extraterminal domain (BET) family proteins such as bromodomain containing 4 (BRD4) are transcription regulators, which localize to cellular enhancer and promoter loci and control the transcription of a wide range of proinflammatory genes. BRD4 regulates transcription by recognizing acetylated histone tail and interacts with transcription factors such as NF-κB p65 and transcription elongation complex [21]. This mechanism facilitates the phosphorylation of RNAPII and therefore promotes transcription initiation and elongation. In the context of proinflammatory gene activation, interaction with NF-κB p65 is especially important, because p65 itself is acetylated at K218, K221, and K310, and the acetylated K310 is recognized by the BRD4 bromodomain to form an active transcription complex [22]. In addition to the specific protein-protein interactions, BRD4 also forms nuclear puncta on super-enhancer genomic regions that exhibit properties of liquid-like condensates [23]. The intrinsically disordered regions (IDRs) of BRD4 are responsible for phase-separated condensates formation with other coactivators on the chromatin loci for transcription elongation [23], which suggests that enrichment of BRD4 at specific enhancers or promoters are key for selection of genes for activation.

Here we demonstrate that a biological effect of prolonged vIL-6 exposures alters the chromatin landscape changes which provides a putative mechanism of overexpression of inflammatory cytokines in vIL-6 pre-exposed cells. We show that an increased amount of BRD4 on chromatin by the prolonged vIL-6 exposure is responsible for transcription deregulation, and repetitive activation of STAT3 appeared to be responsible for the change. BRD4 accumulation prolongs transcription activity at targeted sites with a secondary stimulus by prolonging transcription factor binding and duration of transcriptions. This epigenetic memory may explain the greater risk of chronic inflammatory disease development in KSHV-infected individuals.

## Results

### Biological effects of prolonged vIL-6 exposure on gene transcription and phenotypes

In contrast to temporally regulated hIL-6 expression during wound healing, persistent KSHV infection leads to continuous vIL-6 production from infected cells. Especially, vIL-6 is detected in the serum of KICS patients and the hematopoietic cells in Castleman’s disease patients with a high KSHV viral load [24]. To study the biological consequences of vIL-6 exposure, we first purified recombinant vIL-6 protein from recombinant baculovirus-infected cells and confirmed its biological activity by STAT1 and STAT3 activation [25]. In the study, we showed that longevity and strong phenotypic changes in primary monocytes were induced by vIL-6 exposure for at least 9 days [25]. Here, we prepared for a cell culture model with THP-1 cells which were exposed to vIL-6 continuously for two weeks (we refer to vIL-6/THP-1 hereafter) **(Fig. 1A)**. THP-1 cells, a representative monocyte cell line, was employed because monocytes play a major role in inflammatory responses [26]. Using this cell culture model, we compared the overall transcriptional profiles between vIL-6/THP-1 and parental THP-1 cells followed by secondary stimulation with selected cytokines. We used vIL-6 (100ng/ml) and hIL-6 (100ng/ml), which can activate common downstream signaling pathways, and TGF-β (100ng/ml), which has a non-overlapping pathway, as a secondary stimulation.

**Figure 1.**
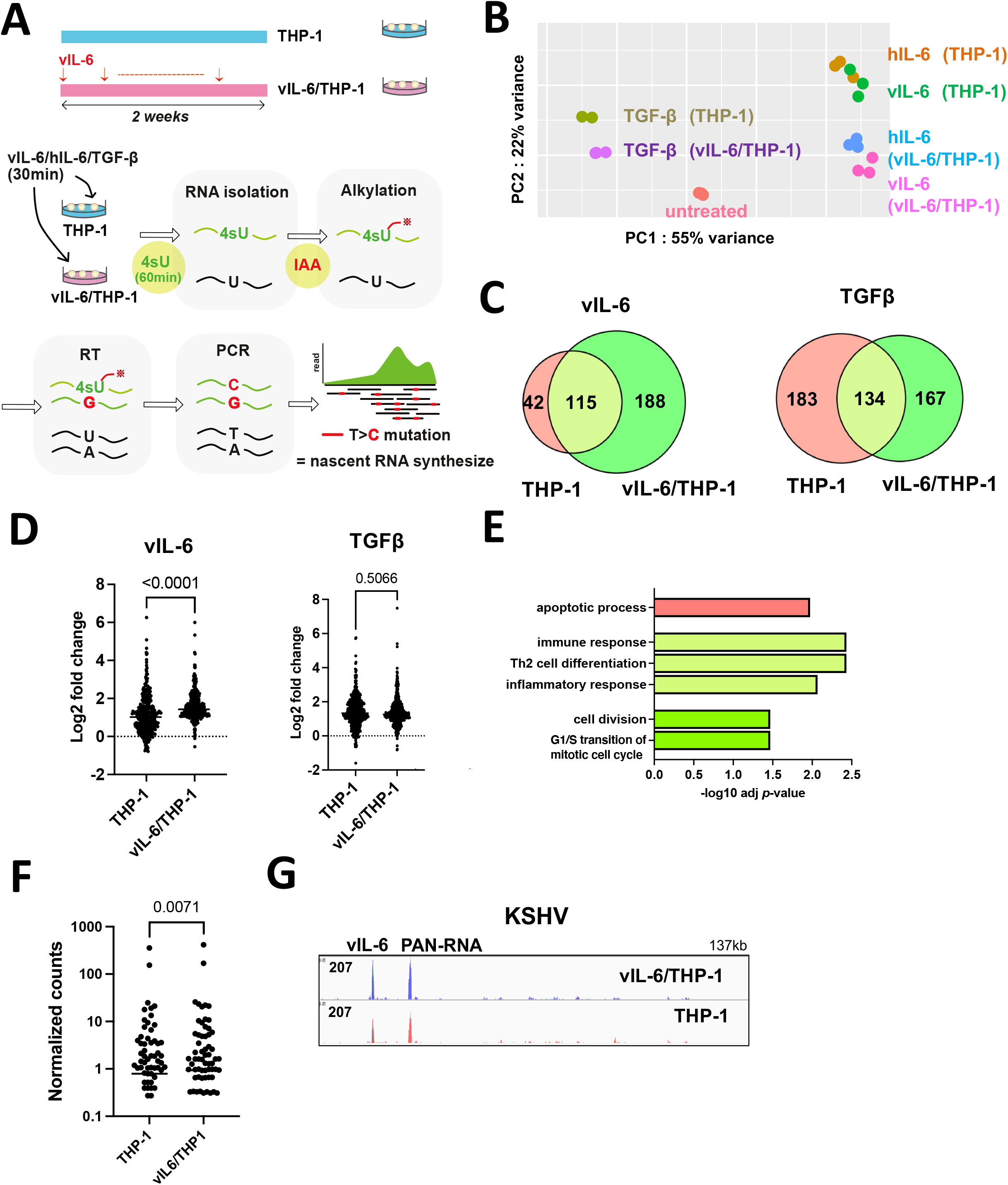
Biological effects of prolonged vIL-6 exposure. **(A)** A schematic diagram of the THP-1 cell culture model. THP-1 cells were treated with or without vIL-6 every other day for 2 weeks (THP-1, vIL-6/THP-1). After withdrawal of the cytokine for 24 hours. vIL-6, hIL-6 and TGF-β were added for 30 minutes for secondary stimulation. Subsequently, 4-Thiouridine (4sU, 300 μM) was added to the culture media and the cells were incubated for 1 hr to label newly synthesized RNA. Newly synthesized RNA was then alkylated by iodoacetamide (IAA, 100mM). Total RNA was isolated for the following analyses. **(B)** Principal component analysis (PCA). Nascent transcribed gene expression in THP-1 and vIL-6/THP-1 cells was shown. vIL-6, hIL-6 and TGF-β were used as secondary stimulation. Untreated THP-1 cells were used as a control. The samples were represented by three biological replicates. The x and y axes show the percentage of variance explained by PC1 (55% variance) and PC2 (22% variance). **(C)** The number of upregulated genes (log2 fold change >1, adj *p*-value < 0.01) after vIL-6 stimulation (left) and TGF-β stimulation (right). Red and green circles represent THP-1 and vIL-6/THP-1 cells, respectively. **(D)** Comparison of up-regulated gene expression between parent THP-1 and vIL-6/THP-1 cells after stimulation with vIL-6 (left, N = 344) or TGFβ (right, N = 484). Data were analyzed using Wilcoxon matched-pairs signed ranked test and shown as median. **(E)** KEGG pathway analysis performed on up-regulated genes (log2 fold change >1, adj *p*-value < 0.01) in either THP-1 or vIL-6/THP-1 cells with vIL-6 stimulation. Each bar represents the pathways that were enriched only in parental THP-1 cells (red), only in vIL-6/THP-1 cells (green), or commonly enriched (light green). Results are presented in descending order of the analysis. **(F)** Comparison of all KSHV gene expression between parent THP-1 and vIL-6/THP-1 cells after KSHV infection (N = 88). Data were analyzed using Wilcoxon matched-pairs signed ranked test and shown as median. **(G)** RNA-sequence reads aligned to the KSHV genome (NC 009333.1) in parent THP-1 (red) and vIL-6/THP-1 cells (blue) after KSHV infection.

To evaluate the newly synthesized RNAs by cytokines stimulation, thiol (SH)-Linked Alkylation for the Metabolic sequencing of RNA (SLAM-seq) was employed [27]. vIL-6/THP-1 and parental THP-1 cells were unstimulated or stimulated by vIL-6, hIL-6 or TGF-β for 30 minutes, followed by incubation with 4-thiouracil (4sU) for 60 minutes to label transcribing RNAs during the incubation periods. In this experimental setting, sequence reads with mutations T>C identify the only nascent synthesized mRNA because 4sU incorporation with chemical alkylation forces incorporation of G instead of A at 4sU positions during the reverse transcription step [27]. Isolated sequence reads with T>C mutation(s) were then compared among control (non-stimulation), vIL-6, hIL-6, and TGF-β **(Fig.1A)**. As shown in **Fig. 1B**, principal component analysis (PCA) demonstrated clearly distinct cellular responses in vIL-6/THP-1 and also among stimuli. In response to vIL-6 stimulation, we observed that the number of upregulated genes (log2 fold change >1, adjusted *p*-value < 0.01) in vIL-6/THP-1 was about twice as many as those with parental THP-1 cells (**Fig. 1C, left**). Similar responses were also seen with hIL-6 as a secondary stimulation (**SFig. 1A**). The vIL-6 and hIL-6 secondary stimulation showed very similar transcriptional profiles (**SFig. 1B**) and downstream pathways (**SFig. 1C**), suggesting that vIL-6 is clearly a functional homolog of hIL-6. On the other hand, the number of upregulated genes with the TGF-β was slightly decreased in vIL-6/THP-1 compared to parental THP-1 cells (**Fig. 1C, right**). In addition, the degree of up-regulated gene expression by vIL-6 or hIL-6 was also increased to a greater extent in vIL-6/THP1 cells compared to parental THP-1 cells (**Fig. 1D, SFig. 1D**), while there was no significant differences with TGF-β (**Fig. 1D**). The results suggested that the enhanced transcription response is a pathway dependent. Gene Ontology (GO) analysis indicated that genes related to immune response, Th2 cell differentiation and inflammatory response were commonly enriched by secondary vIL-6 stimulation in parental and vIL-6/THP-1 cells (**Fig. 1E: light green**), while cell cycle and cell division pathway were additionally activated in vIL-6/THP-1 cells (**Fig. 1E: green**). Consistent with the GO analysis, cell proliferation was increased in vIL-6/THP-1 cells measured by MTT assay (**SFig. 1E**). Despite an increased number of responding genes with the prolonged vIL-6 exposure, the degree of STAT3 activation and gp130 expression (cell surface receptor for vIL-6) was not changed (**SFig. 1F, G**).

Since KICS patients are characterized not only by increased high vIL-6 or hIL-6 but also by increased viral loads, we also examined the effect in KSHV genes expression during de novo infection in vIL-6/THP-1 cells [8]. For this, we infected THP-1 and vIL-6/THP-1 cells with r219.KSHV and performed total RNA-sequencing (RNA-seq). As shown in **Fig. 1F**, overall KSHV gene expression was higher in vIL-6/THP-1 cells compared to parental THP-1 cells. Moreover, vIL-6 and PAN-RNA expression were significantly higher in vIL-6/THP-1 compared to parental THP-1 cells (**Fig. 1G**). Surprisingly, vIL-6 was transcribed as much as PAN RNA in THP-1 cells (**Fig. 1G**). These results suggested that prolonged vIL-6 exposure enhances transcriptional response to secondary vIL-6 and hIL-6 stimulation, and further assists KSHV genes transcription in vIL-6 pre-treated THP-1 cells.

### Prolonged vIL-6 stimulation increased BRD4 occupancies on chromatin

The studies above demonstrated that prolonged vIL-6 exposure increased the number of responding genes followed by vIL-6 or similar (hIL-6) stimuli without altering the amounts of signal-dependent transcription factor (STAT3). These results led us to examine chromatin modification changes induced by prolonged vIL-6 exposure. For this, we first used CHIP-Atlas, a data mining tool for predicting proteins bound to promoters of upregulated genes based on a comprehensive epigenetic database to narrow down possible epigenetic factors. As shown in **Fig. 2A**, STAT1 and STAT3 were commonly extracted as putative responsible transcriptional factors with vIL-6 stimuli both in parental THP-1 and vIL-6/THP-1 cells, demonstrating the validity of this enrichment analysis. Interestingly, transcription coactivators and several transcription factors such as Cyclin dependent kinase 8 (CDK8), Mediator complex subunit 12 (MED12) and Bromodomain-containing protein 4 (BRD4), are likely to be enriched only in the genomic loci in vIL-6/THP-1 cells (**Fig 2A**). BRD4 is a member of the BET (bromodomain and extra terminal domain) family and an important coactivator that binds to H3K27Ac to mediate transcription regulation together with CDK8, MED12 and RNA polymerase II (RNAPII) [28]. Because BRD4 was identified as the most enriched transcription regulator with prolonged vIL-6 exposure (**Fig. 2B**), we next performed Cleavage Under Targets & Release Using Nuclease (CUT&RUN) for BRD4 and RNAPII. In addition, two active histone modification marks (H3K27Ac, H3K4me3) and CTCF were examined to reveal changes in chromatin landscape [29]. Parental THP-1 cells and vIL-6/THP-1 cells were compared in duplicate samples for statistical analyses. Consistent with the CHIP-Atlas results, BRD4 was more enriched in the transcription start sites of up-regulated genes in vIL-6/THP-1 cells (**SFig. 2A**). We also found that BRD4 and H3K27Ac peaks were increased across the genome in vIL-6/THP-1 cells while RNAPII and H3K4me3 were unchanged (**Fig. 2C**). Immunoblotting showed that the total amount of BRD4 modification was not affected with prolonged vIL-6 exposure, as well as H3K27Ac or H3K4me3, demonstrating that increased BRD4 in vIL-6/THP-1 was from the non-chromatin bound fraction **(SFig. 2B)**. HOMER motif analysis for newly accumulated BRD4 genomic regions identified several key monocyte transcription factors such as E74, ETS Transcription Factor 4, and PU.1**SFig. 2C**). PU.1 was also identified as a transcriptional factor likely to be involved in gene activation seen in prolonged vIL-6 exposure by SLAM-seq **(Fig. 2A)**.

**Figure 2.**
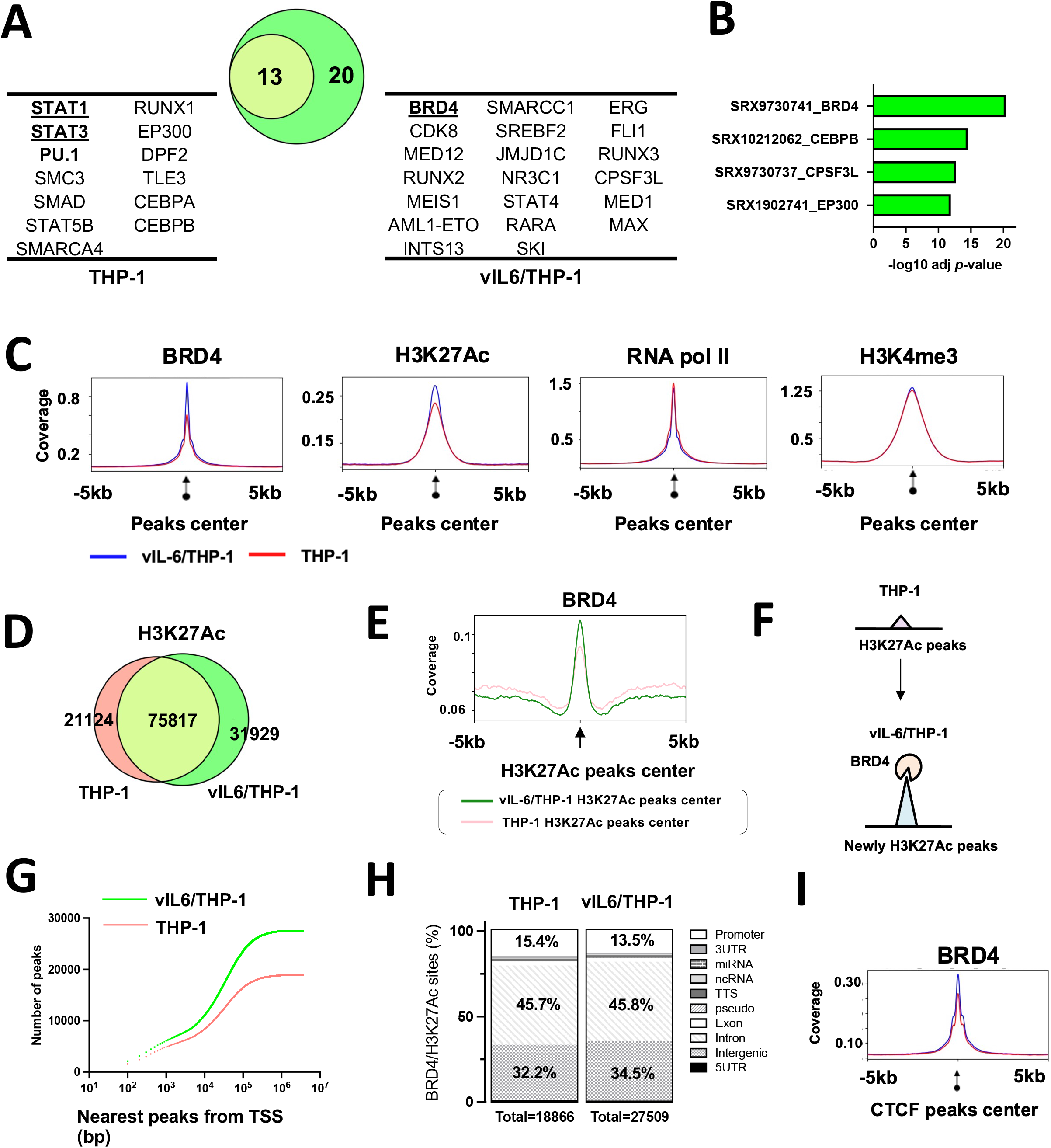
Newly BRD4 recruitment and H3K27Ac translocation in vIl-6/THP-1 cells. **(A)** Enrichment analysis by CHIP-Atlas. Enrichment analysis was performed on the list of genes whose expression were up-regulated by vIL-6 stimulation. The parameters used were as follows; cell type class: blood, threshold for significance: 100, threshold for log10 adjusted *p* value < -10. The Venn diagram shows the relatedness and number of genes up-regulated after vIL-6 stimulation in parental THP-1 (red circle) and vIL-6/THP-1 cells (green circle). **(B)** The predicted transcription factors or mediators selectively enriched only in vIL-6/THP-1 cells. The results were limited to THP-1 cells in Chip-Atlas database. **(C)** CUT&RUN signals in ±5-kbp windows around the center of peaks in vIL-6/THP-1 cells. The list of peaks (p-value < 10-^4^) was extracted using findPeaks (HOMER with default parameters). Enrichment is shown in vIL-6/THP-1 (blue line) and in THP-1 (red line) cells. Images were drawn by plotProfile (HOMER). **(D)** H3K27Ac marks in parental THP-1 and vIL-6/THP-1 cells. The number of H3K27Ac peaks in vIL-6/THP-1 and parental THP-1 cells are depicted by a Venn diagram. **(E)** BRD4 CUT&RUN signals in vIL-6/THP-1 cells. The green line indicates BRD4 peaks±5-kbp around the center of H3K27Ac peaks in vIL-6/THP1 cells while the pink line indicates those in parental THP-1 cells. **(F)** Schematic model of H3K27Ac translocation and BRD4 recruitment at newly emerged H3K27Ac regions in vIL-6/THP-1 cells. **(G)** Distances of the nearest TSS to overlapping peaks of BRD4 binding and H3K27Ac enrichment sites. Total peaks counts are shown. *P* values were calculated by the Kolmogorov–Smirnov test. **(H)** Annotation of overlapping peaks between BRD4 and H3K27Ac enrichment sites. Each annotation and its proportion were calculated by annotatePeaks.pl (Homer; parameters; default). Promoter regions are defined as ± 1-kbp from TSS. **(I)** BRD4 CUT&RUN signals in ±5-kbp windows around the center of CTCF peaks in vIL-6/THP-1 cells (blue) and parental THP-1 cells (pink).

We next examined the position and association of the newly accumulated BRD4 with H3K27Ac to reveal the changes of active genomic hubs. The CUT&RUN studies identified 31,929 new H3K27Ac regions, while 21,124 regions found in parental cells were no longer modified by H3K27Ac in vIL-6/THP-1 cells **(Fig. 2D).** As shown in **Fig. 2E**, BRD4 was enriched more significantly and sharply at the centers of H3K27Ac peaks in vIL-6/THP-1 cells. The genomic regions, where BRD4 and H3K27Ac peaks overlap, were also increased from 18,886 to 27,509 by prolonged vIL-6 exposure (**SFig. 2D**), suggesting that newly accumulated BRD4 was recruited in newly established H3K27Ac loci in vIL-6/THP-1 cells (**Fig. 2F**). RNAPII and H3K4me3 were also enriched in the genomic regions, where BRD4 and H3K27Ac peaks overlapped (**SFig. 2E**). Closer examination of these enriched sites demonstrated that while the overall number of BRD4/H3K27Ac overlapped peaks around TSS was increased (**Fig. 2G**), the relative abundance of the peaks at the promoter region decreased from 15.4% to 13.5 % in vIL-6/THP-1 cells (**Fig. 2H**). These results may suggest that a larger fraction of newly generated active hubs is located at intragenic regions, perhaps at newly activated enhancers. The CUT&RUN results with the CTCF antibody also suggested that prolonged vIL-6 stimulation increased overall co-occupancies between CTCF and BRD4 across the genome (**Fig. 2I**), suggesting the establishment of activated chromatin hubs in vIL-6/THP-1 cells. The new BRD4 and H3K27Ac peaks were co-localized to promoter regions of genes associated with cellular response to DNA damage stimulus and cell division, such as cyclin E1, which is consistent with the results from the SLAM-seq analysis **(SFig. 2F, G)**.

### Enhanced response to LPS via prolonged vIL-6 stimulation

Synergistic interactions between NF-κB and STAT3 induce the hyperactivation of NF-κB followed by the production of various inflammatory cytokines such as hIL-6 [18]. KSHV-infected cells are known to have elevated NF-κB activation and STAT pathways are constitutively activated in primary effusion lymphomas [30, 31]. Accordingly, we examined if repetitive STAT activation by vIL-6 exposure alters NF-κB mediated cell responses in *cis.* To study this, we stimulated vIL-6/THP-1 cells with LPS, a strong inducer of NF-κB pathway activation, and examined the cellular response. We first confirmed that 100 ng/ml of LPS stimuli for 6 hours is sufficient for detecting hIL-6 production by ELISA assay **(SFig. 3A)**, and selected this as a time point to collect samples in order to minimize the secondary effects from the newly produced cytokines with LPS stimuli. We measured a series of downstream inflammatory cytokines produced with Olink, a multiplex proximity extension assay (PEA) [32]. The results showed that LPS upregulated the expression of 21/41 inflammatory cytokines (log2 fold change >1) in parental THP-1 cells (**Fig. 3A**). Among them, the expression of 15 cytokines, such as hIL-6, IL-10, IL-1β, and CCL8, was enhanced in vIL-6/THP-1 cells compared to parental THP-1 cells (**Fig. 3A, B**). IFNα stimulation also induced more cytokines in vIL-6/THP-1 cells similarly to that of LPS stimulation (**SFig. 3B, C**), suggesting that prolonged vIL-6 exposure to monocytes sensitized cells to secondary stimuli, reinforcing the notion that enhanced activation is due to the formation of active chromatin complexes at local chromatin.

**Figure 3.**
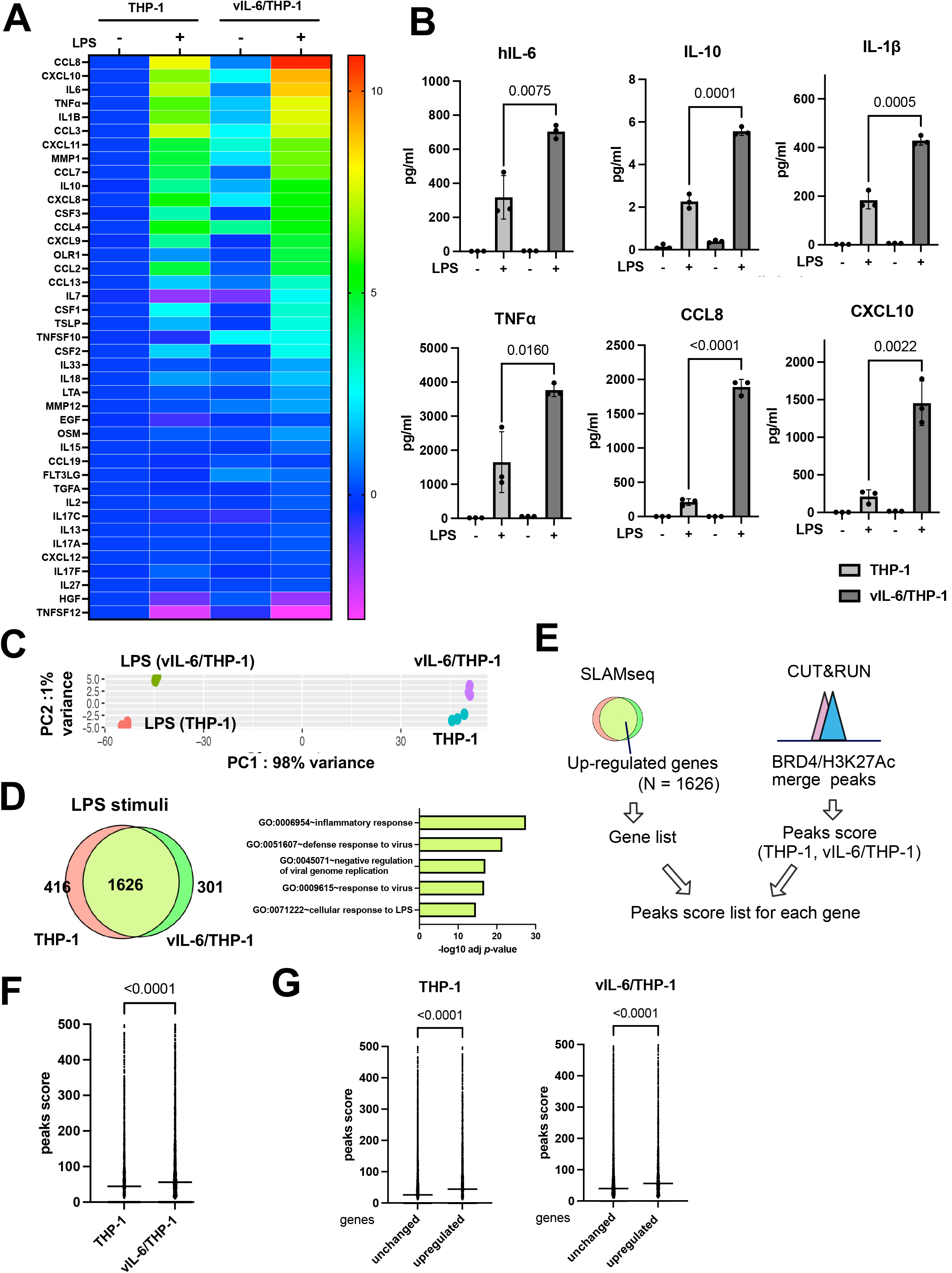
Prolonged vIL-6 exposure enhances inflammatory response to LPS through the novel accumulation of BRD4 and H3K27Ac. **(A)** Heatmap showing the results of Olink® Target 48 Cytokine panel analysis. LPS (100ng/ml) was added to parental THP-1 or vIL-6/THP-1 cells for 6 hours. Cytokine production in untreated THP-1 cells was set as 1 and log_2_ fold activation relative to untreated THP-1 cells is shown. Samples were prepared in triplicate and the mean value were shown. **(B)** Individual inflammatory cytokine production determined by Olink proximity extension assay. Data was analyzed using two-sided unpaired Student’s *t* test and shown as mean ± SD. **(C)** PCA based on nascent transcribed gene expression in THP-1 cells and vIL-6/THP-1 cells after LPS stimulation. The samples were represented by 3 biological replicates. The x and y axes show the percentage of variance explained by PC1 (98% variance) and PC2 (1% variance). **(D)** The number of up-regulated genes (log2 fold change >1, adj *p*-value < 0.01) in THP-1 and vIL-6/THP-1 cells after LPS stimulation (left) and KEGG pathway analysis performed on commonly up-regulated genes (right). **(E)** Schematic diagram for comparing peaks score of up-regulated genes between THP-1 and vIL-6/THP-1 cells. Commonly up-regulated genes by LPS stimulation were obtained from SLAMseq data and BRD4/H3K27Ac merge peaks score were obtained from CUT&RUN data. The peak scores were associated with each up-regulated gene. **(F)** BRD4/H3K27Ac merge peak score for common upregulated genes in THP-1 and vIL-6/THP-1 cells (N = 1626). Data were analyzed using Wilcoxon matched-pairs signed ranked test and shown as median. **(G)** BRD4/H3K27Ac peak score between up-regulated genes (N = 1626) and unchanged genes (log2 fold change >1 and < -1, adj *p*-value > 0.01) (N = 4582) in THP-1 (left) and vIL-6/THP-1 cells (right). Data were analyzed using Mann-Whitney test and shown as median.

The transcription profile in response to LPS was also accompanied by SLAMseq analysis. LPS (100ng/ml) and 4sU were added to parental THP-1 cells or vIL-6/THP-1 cells at the same time and incubated for 6 hours to label nascent transcribed RNAs. PCA demonstrated that LPS strongly changed the gene transcription profile, and the variance between THP-1 and vIL-6/THP-1 cells was further increased by LPS stimulation (**Fig. 3C**). As expected, the inflammatory response was identified as the most enriched pathway in commonly up-regulated genes (N = 1626) (**Fig. 3D**). Next, we investigated whether the increased BRD4 and H3K27Ac occupancies in vIL-6/THP-1 cells contributed to the increased inflammatory gene transcription by LPS stimulation. Commonly up-regulated genes were extracted and the peak scores of the BRD4 and H3K27Ac overlapping regions in THP-1 and vIL-6/THP-1 cells were compared (**Fig. 3E**). As shown in **Fig. 3F**, co-occupancy of BRD4 and H3K27Ac was indeed associated with increased transcription activation with LPS. The increased responses to LPS stimuli were also associated with an increased BRD4 and H3K27Ac co-occupancy compared to genes whose expression was not changed (**Fig. 3G**). These results suggested that prolonged vIL-6 exposure enhances the inflammatory response via increased acetylation of H3K27 at BRD4 recruited sites.

### BRD4 is responsible for prolonging the hIL-6 transcription burst with LPS

Because BRD4 was enriched in the promoter regions of hIL-6, hIL-10 and IL-1β with the prolonged vIL-6 exposure **(SFig. 4A)** and are associated with increased production of cytokines in culture media **(Fig. 3A)**, we next studied how BRD4 recruitment contributes to increasing transcripts. We speculated that increased occupancies of BRD4, which contains an intrinsically disordered domain to form liquid-liquid phase separation (LLPS) [33], may increase the recruitment and duration of p65 binding at promoter regions. This mechanism may extend the duration or robustness of transcription after stimulation. Accordingly, we examined the recruitment of p65 after LPS stimuli in vIL6/THP-1 cells and compare it with parental THP-1 cells. The proximity ligation assay showed that the association between BRD4 and p65 was increased after LPS stimulation in vIL6/THP-1 cells (**Fig. 4A**), while the total p65 amount in the nucleus after LPS stimulation was not significantly changed in vIL-6/THP-1 cells (**Fig. 4B**). Chip-qPCR also showed increased p65 recruitment on the hIL-6, IL10, and IL-1β promoters by LPS stimuli in vIL6/THP-cells (**Fig. 4C**). Transcription frequencies were also determined by qRT-PCR analysis of nascent RNA **(SFig. 4B)**. As shown in **Fig. 4D**, hIL-6, IL-10 and IL-1β transcripts were produced more rapidly (hIL-6 and IL-10) and/or continuously (IL-1β) in vIL-6/THP-1 cells after stimulation. Importantly, BRD4 occupancies are responsible for the increased transcription, because OTX-015, a BRD4 inhibitor, drastically suppressed the transcription in both parental THP-1 and vIL-6/THP-1 cells, and the effect was more pronounced in vIL-6/THP-1 cells (**Fig. 4E**). Taken together, prolonged vIL-6 exposure changed the chromatin landscape to be favorable to the response of inflammatory secondary stimuli, and establishment of active chromatin hubs with BRD4 recruitment is critical for this response (**Fig. 5**).

**Figure 4.**
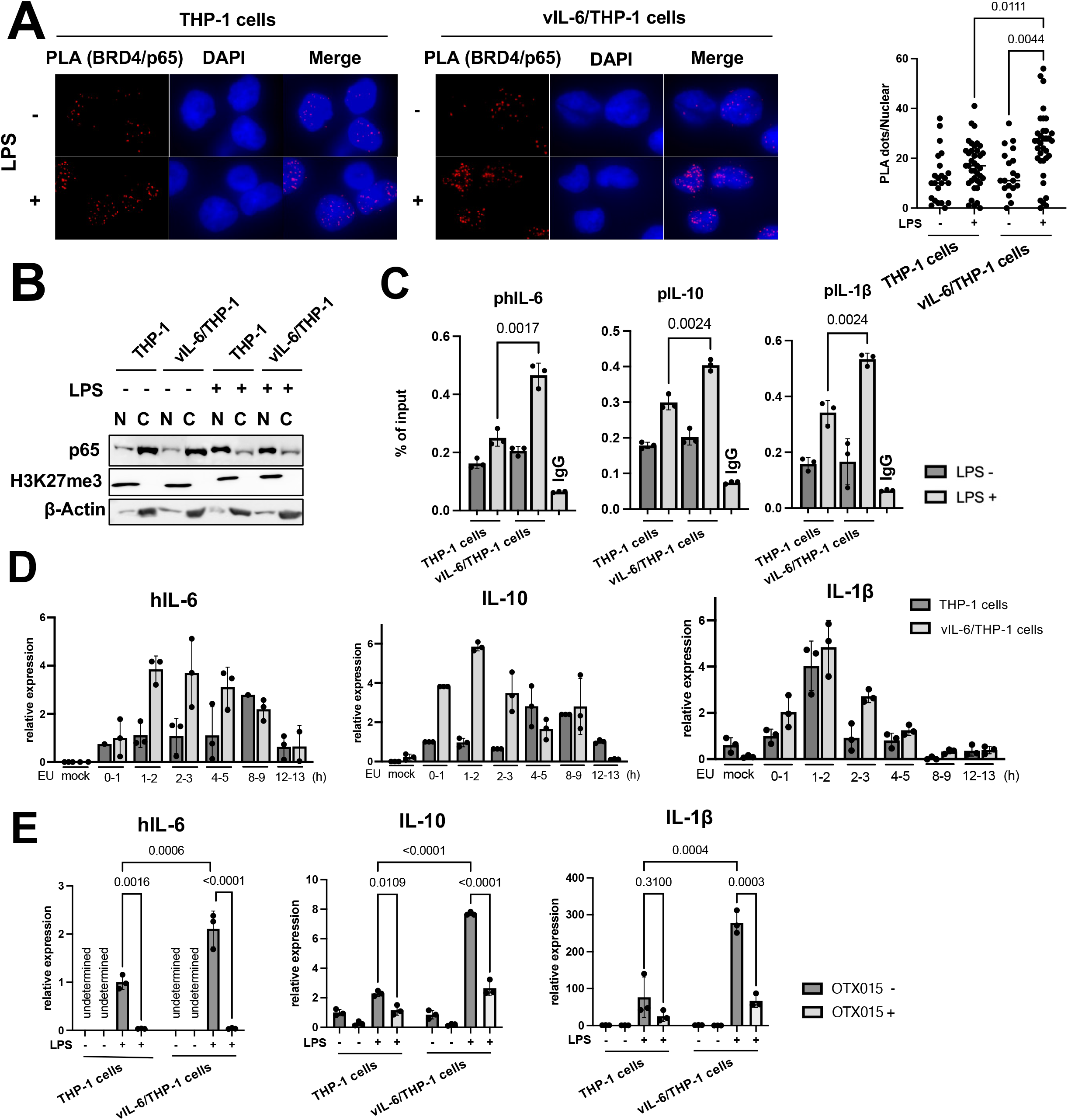
Association of BRD4 and p65 is responsible for prolonging hIL-6 transcription burst by LPS. **(A)** Proximity extension assay (PLA) to visualize BRD4 and p65 interaction. THP-1 and vIL-6/THP-1 cells were incubated with LPS (100ng/ml) for 6 hours, fixed by 4% PFA/PBS for 15min, permeabilized by 0.15% TritonX/PBS for 10min and PLA was performed. Representative images of PLA (red dots) and quantification of PLA are shown (right). Data were analyzed using a one-way ANOVA test. **(B)** Immunoblotting of p65, H3K27me3 and β-actin in the nuclear and cytoplasmic fractions. THP-1 and vIL-6/THP-1 cells were incubated with LPS (100ng/ml) for 6 hours and cell fractionation was performed. **(C)** The levels of p65 enrichment in the hIL-6, hIL-10 and IL-1β promoter regions were detected using ChIP-qPCR. Normal Rabbit IgG antibody was used as negative control. Data was analyzed using two-sided unpaired Student’s *t* test and shown as mean ± SD. **(D)** Nascent RNA expression in THP-1 and vIL-6/THP-1 cells after LPS stimulation. Cells were incubated with LPS and ethynyl uridine (EU) was added 0h, 1h 2h, 4h, 8h, 12h after LPS stimulation for 1 hour. Mock samples were prepared without EU for 1 hour incubation with LPS. **(E)** The level of hIL-6, IL-10 and IL-1β gene expression in response to OTX-015 treatment. THP-1 and vIL-6/THP-1 cells were incubated with OTX-015 (10nM) for 4 hours followed by LPS (100ng/ml) for 1 hour. RNA was then collected and transcribed into cDNA for RT-qPCR. 18S rRNA expression was used for internal control. Data were analyzed using a one-way ANOVA test.

**Figure. 5.**
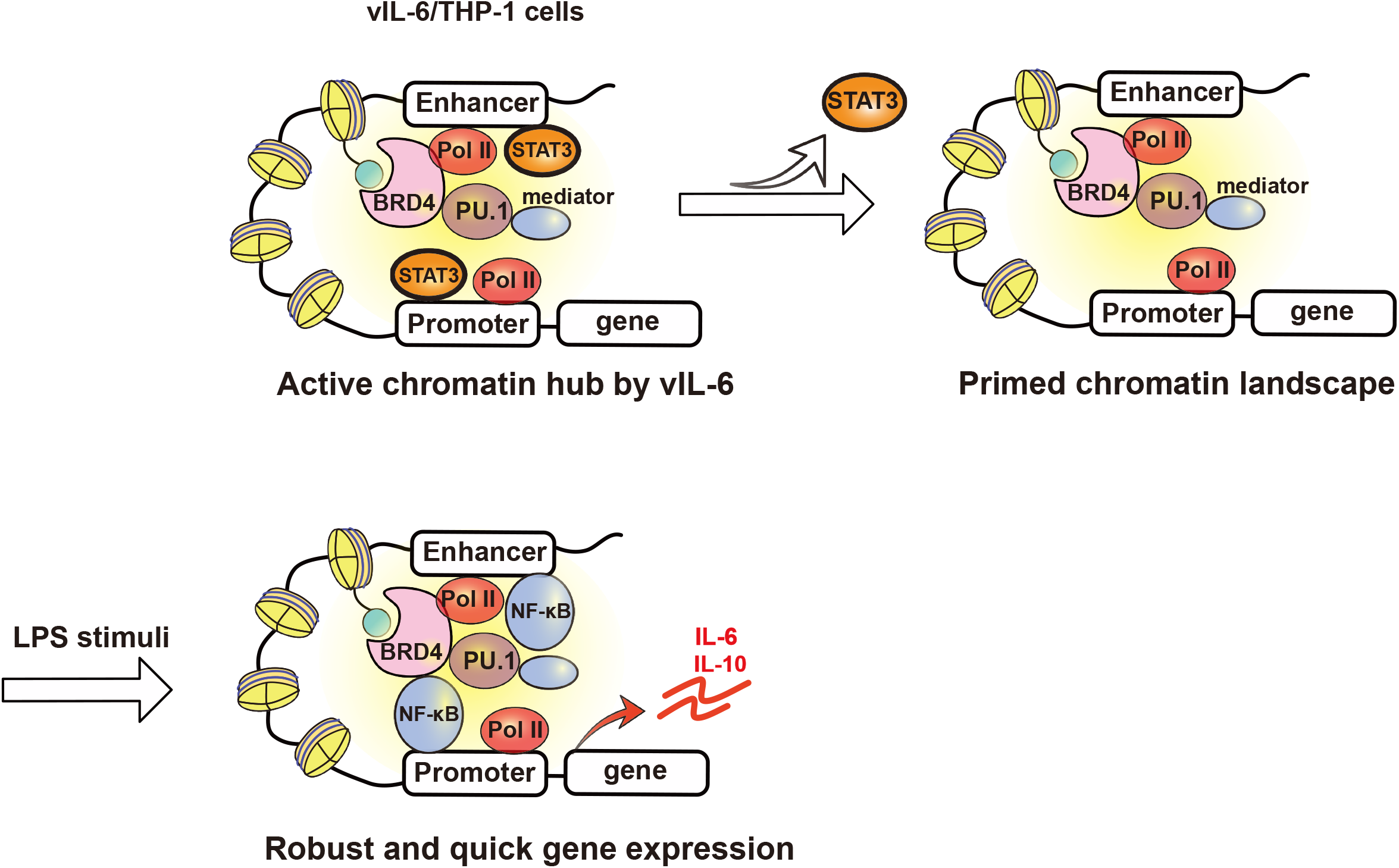
Schematic model of transcription reprogramming by prolonged vIL-6 exposure. Signal-dependent transcription factors such as STAT3 are recruited at up-regulated gene promoters by vIL-6, and BRD4 and H3K27Ac are increasingly co-occupied at those promoters by prolonged vIL-6 exposure. After the removal of the stimulus, STAT3 is removed from the promoters; however, the activated H3K27Ac modification and BRD4 remained and established a “primed” chromatin state for subsequent activation. Subsequent LPS stimulation activated hIL-6 or IL-10 genes that had been primed by vIL-6, more quickly and robustly.

## Discussion

In this study, we applied SLAM-seq for most of the RNA-seq experiments to increase the resolution to identify direct target genes (**Fig. 1, 3C**). A short window after the respective stimuli was utilized to examine robustness and differences made in direct target gene expression with prolonged vIL-6 stimuli. By measuring nascent RNA that was labelled shortly after stimulation, we also avoid indirect, or secondary activation. The study clearly demonstrated that direct target genes are significantly overlapped with that of hIL-6 (**Fig.1B**). This also suggests that these two signaling ligands are interchangeable, making it easier to have a continuously stimulated IL-6 signaling pathway in KSHV infected individuals.

Cytokines that activate STAT3 (e.g. hIL-6) and NF-κB (e.g. TNF-α) synergistically increase the production of various inflammatory cytokines, called IL6-Amp. Since vIL-6 has the same biological activity as hIL-6 **(Fig.1B, SFig. 1B, C),** it is reasonable to speculate that vIL-6 bypasses hIL-6 expression to trigger the IL6-Amp in vivo. The vIL-6 can bind to gp130 alone and does not require the high affinity IL-6 receptor [34, 35]. This is in clear contrast to hIL-6, which must bind to its high affinity receptor before a signaling complex can be formed with its signal transducer, gp130 [36, 37]. Importantly, the gp130 is widely expressed in human tissues, and the notion that vIL-6 activates downstream JAK/STAT pathway without requiring binding to the classical gp80 receptor [38–41] suggests that vIL-6 would initiate signaling in a wider variety of cell types [40, 42]. In addition, previous studies by Dr. Nicholas’s group further demonstrated that vIL-6 can function though intracellularly via activation of gp130 at the endoplasmic reticulum in infected cells [43, 44]. The study suggested that vIL-6 may support the growth and survival of KSHV-infected cells in an autocrine manner and such unique signaling activities contribute to the maintenance of latently infected cells without alarming bystander cells. Our recent study with primary monocyte infection as well as this study agreed very well with the observation, in which prolonged vIL-6 stimuli shifted the cell transcription program towards cell growth and anti-apoptotic signaling (**Fig. 1E, SFig. 1E**). More importantly, similar phenotypes were also seen in wild-type KSHV primary infection but not infection with vIL-6 stop virus [25].

Prolonged vIL-6 stimuli similarly increased the number of responding genes to secondary vIL-6 and hIL-6 stimulation whereas TGF-β did not show enhanced responses (**Fig. 1C, D**). Generally, chronic inflammation leads to a stronger inflammatory response, which is described as immunological training [45]. In immunological training, the primary stimulus will prime specific regulatory elements in enhancer regions. This established chromatin landscape allows faster and increased transcriptional activation of the secondary stimulus, leading to an enhanced inflammatory response. At the chromatin level, immunological training has been shown to induce long-lasting epigenetic remodeling of regulatory elements [46]. This remodeling includes changes in activating histone marks such as H3K27Ac and H3K4me3 [46, 47]. In this study, signal-dependent transcription factors such as STAT1 and STAT3 motifs were enriched at up-regulated gene promoters, and BRD4 and H3K27Ac were increasingly co-occupied at those promoters **(Fig. 2C)**. After removal of the stimulus, STAT3 is likely to be left from the promoters; however, our study suggested that BRD4 and the activated H3K27Ac modification remained and established a “primed” chromatin state for subsequent activation **(Fig. 5)**. As a result, LPS activated selected genes that had been primed by STAT3, more quickly and strongly **(Fig. 4E)**. Indeed, cellular enzymes that are known to be enriched at enhancers such as PU.1, MED1, SWI/SNF complexes are predicted to be co-occupied at newly responding gene promoters **(Fig. 2A, SFig. 2C)**. These observations support that vIL-6 stimulation induced transcription reprogramming, which favors a subsequent inflammation response.

What would be the evolutional advantage for maintaining vIL-6 in KSHV genome? Interestingly, vIL-6 is reported to be constitutively expressed in some cell populations even during latent infection [13, 48]. We think that immunological training effects are utilized for the establishment of active latent chromatin for inducible lytic gene expression. Consistent with this hypothesis, BRD4, H3K27Ac, and H3K4me3 are all highly accumulated at the KSHV ori-Lyt region in latent chromatin [49]. We think that KSHV is likely to utilize host defense mechanisms to recruit BRD4 on its genome to establish a “primed” latent chromatin for further reactivation, which is partly seen in increased KSHV transcription in vIL-6/THP-1 cells (**Fig. 1F, G**). Perhaps, related to this, we recently showed that the vIL-6KO virus could not reprogram primary infected monocytes and could not replicate continuously [25]. These cellular and virological vIL-6 functions are likely to contribute to the highly inflammatory phenotypes seen in KSHV-infected patients.

In summary, we reported that long-term vIL-6 exposure increased BRD4 and H3K27Ac co-occupancies at inflammatory-related genes, resulting in enhanced inflammatory responses to LPS. BRD4 is clearly the center of the altered cell responses caused by prolonged vIL-6 exposure, in which the pre-located BRD4 on the promoter enabled transcription factors and cofactors that are brought in proximity by p65 to interact more rapidly and steadily upon LPS stimulation. Changes in the enhancer-promoter genomic interactions by long-term exposure to vIL-6 remain a promising avenue for further research. Moreover, animal models with continuous vIL-6 exposure should reveal how vIL-6 trained memory polarizes immune cell communications and react to secondary stimuli, which will provide valuable information for a better understanding the pathogenesis of KICS. Although continued treatment with BET inhibitors is known to cause large genomic structural changes and rapidly induce drug-resistant cells [50, 51], short-term treatment with well-scheduled drug holidays may be beneficial to reverse the immunological training memory established by KSHV vIL-6.

## MATERIALS AND METHODS

### Chemicals, reagents and antibodies

Dulbecco’s modified minimal essential medium (DMEM), fetal bovine serum (FBS), phosphate-buffered saline (PBS), Trypsin-EDTA solution, 100 X penicillin–streptomycin– L-glutamine solution were purchased from Thermo Fisher (Waltham, MA). Puromycin and G418 solution were obtained from InvivoGen (San Diego, CA). Hygromycin B solution was purchased from Enzo Life Science (Farmingdale, NY).

The following antibodies were used for CUT & RUN, immunoblotting and flow cytometry: rabbit anti-BRD4 (Cell Signaling, E2A7X), rabbit anti-RNAPII (Millipore, clone CTD4H8), rabbit anti-H3K27ac (CST, clone D5E4), rabbit anti-H3K4me3 (Cell Signaling, clone C42D8), mouse anti-β-actin (Santa Cruz 47778), rabbit anti-gp130 (CST), rabbit IgG (CST, clone DA1E). Rabbit anti-vIL-6 was a gift from Dr. Robert Yarchoan (NIH/NCI).

### Cells

THP-1 cell lines was obtained from ATCC. Cells were maintained in an RPMI 1640 medium supplemented with 1% fetal bovine serum (FBS; Invitrogen), 2 mM L-glutamine and 100 U/ml penicillin and 100 μg/ml streptomycin at 37°C in a humidified 5% CO_2_. THP-1 cells were passaged two to three times per week when reaching 1 x 10^6^ cells/ml.

### Purification of recombinant protein

*Spodoptera frugiperda* Sf9 cells (Millipore) were maintained in Ex-Cell 420 medium (Sigma), and recombinant baculoviruses were generated with the BAC-to-Bac system as previously described [52] [53].

### SLAM-seq

SLAM-seq [27] was performed using the SLAMseq Kinetics Kit (Lexogen GmbH, Vienna, Austria) according to the manufacturer’s standard protocol.

Briefly, biological replicate cultures of THP-1 cells were incubated with vIL-6 (100µg/ml) for 30 min or 2 weeks. Subsequently, 4-Thiouridine (s4U; 300 μM) was added to the culture media and the cells were incubated for 1 h in order to label newly synthesized RNA. As for the LPS (100ng/ml) stimulation, s4U was added to the culture media simultaneously with LPS, and cells were incubated for 6 hours. Total RNA was isolated and then the 4-thiol groups in the s4Uracil-labeled transcripts were alkylated with iodoacetamide (IAA). QuantSeq 3′ mRNA-Seq (FWD) (Lexogen, Inc.) Illumina-compatible, indexed sequencing libraries were prepared from alkylated RNA samples (100 ng) according to the manufacturer’s protocol for oligo(dT)-primed first-strand cDNA synthesis, random-primed second-strand synthesis, and library amplification. Libraries were multiplex sequenced (1 × 100 bp, single read) on an Illumina HiSeq 4000 sequencing system. SLAM-Seq datasets were analyzed using the T > C conversion-aware SLAM-DUNK (Digital Unmasking of Nucleotide conversion-containing k-mers) pipeline utilizing the default parameters [27, 54]. Briefly, nucleotide conversion-aware read mapping of adapter-and poly(A)-trimmed sequences to the human GRCh38/hg38 reference genome assembly was performed with NextGenMap [55]. Alignments were filtered for those with a minimum identity of 95% and minimum of 50% of the read bases mapped. For multi-mappers, ambiguous reads and non-3′ UTR alignments were discarded, while one read was randomly selected from multimappers aligned to the same 3′ UTR. SNP calling (coverage cut-off of 10X and variant fraction cut-off of 0.8) with VarScan2 [56] was performed in order to mask actual T > C SNPs. Non-SNP T > C conversion events were then counted and the fraction of labeled transcripts was determined. All results were used for downstream analyses, such as nascent transcript analysis and differential expression analysis (DESeq2) [57].

### Bioinformatics analysis of SLAM-seq data

The UCSC Genome Browser was used to convert RefSeq IDs to gene symbols (refGene). The resulting data were first filtered by log2 fold change and sorted by adjusted p-value from lowest to highest. The resulting up-regulated genes (log2 fold change >1 and adjusted *p*-value <0.01) were extracted. For functional enrichment analysis (KEGG pathway analysis), the official gene symbols were applied in DAVID web service tools [58]. For transcription factor analysis, gene symbols were submitted to ChIP-Atlas [59] to analyze common regulators and to predict transcription factor binding. The following parameters were used; Organism (annotation): Homo sapiens (hg38), Experiment type; TFs and others, Cell type Class; Blood, Threshold; 100. The output factors were then listed and plotted in a Venn diagram.

### RNA-sequencing

vIL-6/THP-1 and parental THP-1 cells were infected with or without r219.KSHV for 72h and RNA was purified using the Quick-RNA miniprep kit (Zymo Research, Irvine, CA, USA). Indexed, stranded mRNA-seq libraries were prepared from total RNA (100 ng) using the KAPA Stranded mRNA-Seq kit (Roche) according to the manufacturer’s standard protocol. Libraries were pooled and multiplex sequenced on an Illumina NovaSeq 6000 system (150-bp, paired-end, >30 × 106 reads per sample). RNA-Seq data was analyzed using a Salmon-tximport-DESeq2 pipeline. Raw sequence reads (FASTQ format) were mapped to the reference human genome assembly (GRCh38/hg38, GENCODE release 36) and quantified with Salmon [60]. Gene-level counts were imported with tximport [61] and differential expression analysis was performed by DESeq2 [57], which are visualized by Venn diagram.

### Cleavage under targets and release using nuclease (CUT&RUN)

CUT&RUN [29] was performed essentially by following the online protocol developed by Dr. Henikoff’s lab with a few modifications to fit our needs. Cells were washed with PBS and wash buffer [20 mM HEPES-KOH pH 7.5, 150 mM NaCl, 0.5 mM Spermidine (Sigma, S2626), and proteinase inhibitor (Roche)]. After removing the wash buffer, cells were captured on magnetic concanavalin A (ConA) beads (Polysciences, PA, USA) in the presence of CaCl2. Beads/cells complexes were washed three times with digitonin wash buffer (0.02% digitonin, 20 mM HEPES-KOH pH 7.5, 150 mM NaCl, 0.5 mM Spermidine and 1x proteinase inhibitor), aliquoted, and incubated with specific antibodies (1:50) in 250 μL volume at 4℃ overnight. After incubation, unbound antibody was removed by washing with digitonin wash buffer three times. Beads were then incubated with recombinant Protein A/G–Micrococcal Nuclease (pAG-MNase), which was purified from E.coli in 250 μl digitonin wash buffer at 1.0 μg/mL final concentration for 1 h at 4 °C with rotation. Unbound pAG-MNase was removed by washing with digitonin wash buffer three times. Pre-chilled digitonin wash buffer containing 2 mM CaCl_2_ (200 μL) was added to the beads and incubated on ice for 30 min. The pAG-MNase digestion was halted by the addition of 200 μl 2× STOP solution (340 mM NaCl, 20 mM EDTA, 4 mM EGTA, 50 μg/ml RNase A, 50 μg/ml glycogen). The beads were incubated with shaking at 37 °C for 10 min in a tube shaker at 300 rpm to release digested DNA fragments from the insoluble nuclear chromatin. The supernatant was then collected by removing the magnetic beads. DNA in the supernatant was purified using the NucleoSpin Gel & PCR kit (Takara Bio, Kusatsu, Shiga, Japan). Sequencing libraries were then prepared from 3 ng DNA with the Kapa HyperPrep Kit (Roche) according to the manufacturer’s standard protocol. Libraries were multiplex sequenced (2 × 150 bp, paired-end) on an Illumina NovaSeq 6000 system to yield ∼15 million mapped reads per sample. With separate replicated experiments, qPCR was used to examine enrichment at selected genomic regions. Primer sequences are provided in **Table.S1.**

CUT&RUN sequence reads were processed with fastp [62] and aligned to the human GRCh38/hg38 reference genome assembly with Bowtie2 and yielding mapped reads in BAM files [63]. Hypergeometric Optimization of Motif EnRichment (HOMER) v4.11 was used for peak detection and their annotation utilizing the defalt parameters described in the developer’s manual [31]. Deeptools was used for making plotprofiles utilizing the defalt parameters [64]. Peaks and read alignments were visualized using the Integrated Genome Browser [65].

### Olink analysis

1.5 × 10^6^ THP-1 cells or vIL-6/THP-1 cells were prepared in triplicate and washed twice by PBS10ml and then suspended with 1ml fresh RPMI medium in 12 well plates. Inflammatory cytokines such as LPS (100ng/ml), IFNα(100ng/ml), vIL-6 (100ng/ml) and hIL-6 (100ng/ml) were then added to each well and cultured at 37℃, 5% CO_2_. 6 hours after incubation, supernatants were harvested and centrifuged (3,000 rpm, 3min) to remove the remaining cells. The resulting supernatant samples were stored at -20℃ for approximately two weeks until they were sent for analysis. Samples wrapped in dry ice were then sent to Olink Proteomics (Temple City, CA). Olink’s Proximity Extension Assay (PEA) technology uses antibody pairs conjugated to unique oligonucleotides and is quantified via PCR. When both antibodies of a pair bind the target protein simultaneously, their respective conjugated oligonucleotides are brought into proximity, facilitating hybridization. The oligonucleotide sequence is then extended by DNA polymerase, amplified, and measured by qPCR to determine the sample’s initial protein abundance. Raw analyte expression values after PCR underwent multiple rounds of transformation by Olink, including a log2 transformation, and were returned as normalized protein expression (NPX) values [66]. For evaluating inflammatory cytokine production in the culture medium, the Olink® Target 48 Cytokine panel was selected by multiple investigators. The data was processed with Olink Insight Stat Analysis software.

### RT-qPCR

Total RNA was extracted using the Quick-RNA miniprep kit (Zymo Research, Irvine, CA, USA). A total of 1 μg of RNA was incubated with DNase I for 15min and reverse transcribed with the High Capacity cDNA Reverse Transcription Kit (Thermo Fisher, Waltham, MA USA). The resulting cDNA was used for qPCR. 10 ml SYBR Green Universal master mix (Bio-Rad) was used for qPCR according to the manufacturer’s instructions. Each sample was normalized to 18S ribosomal RNA, and the ddCt fold change method was used to calculate relative quantification. All reactions were run in triplicate. Primer sequences used for qRT-PCR are provided in the **Table. S1.**

### Immunoblotting

Cells were washed with PBS, lysed in lysis buffer (50 mM Tris-HCl [pH 6.8], 2% SDS, 10% glycerol) and boiled for 3min. The protein concentrations of the lysates were quantified with a BCA Protein Assay Kit (Thermo Fisher). Protein samples were separated by SDS-PAGE using 10% agarose gel and transferred to transfer membranes (Millipore-Sigma, St. Louis, MO, USA), which were incubated in 5% nonfat milk (A600669, Sangon) at room temperature for 2 hours. The membrane was incubated with the primary antibody at 4°C overnight or at room temperature for 2 hours. The membrane was then incubated with horseradish-peroxidase-conjugated secondary antibody or Alexa-647-conjugated secondary antibody at 25°C for 1 hour. For cell fractionation, cells were suspended with hypotonic buffer (20mM Tris-HCl(pH 7.4), 10mM NaCl, 3mM MgCl2, 0.5mM DTT and proteinase inhibitor cocktail) for 15min on ice and 0.5% NP-40 was added. Cells were then centrifuged for 10 minutes at 3000rpm to collect the supernatants (cytoplasm fraction). Pellets were washed with PBS 2 times and suspended with protein lysis buffer (nuclear fraction).

### Flow cytometry

Cells in culture were washed twice with PBS and resuspended in FACS buffer (PBS supplemented with 1% FBS) at 1×10^6^ cells/ml. Cells were then stained for 2 hours at room temperature with the gp130 antibodies (1:1000) followed by Alexa-647-conjugated secondary antibody for 1 hour. For intracellular nuclear staining to measure cell cycle, cells were washed twice with PBS and incubated with propidium iodide (PI) solution (50ng/ml PI, 0.2% NP-40, 0,25mg/ml RNase) for 15 min at 4°C and then for 30 min at 37°C in the dark. Flow cytometry was carried out by using a BD Acuri instrument (BD Biosciences) and data analysis was performed using FlowJo v10.8.1 (Tree Star) by gating on live cells based on forward versus side scatter profiles. Cell cycle was calculated by Watson model implemented in FlowJo according to the manufacture’s protocol.

### Capture of nascent RNAs

To capture nascent RNAs, 0.4 mM EdU was added into the culture medium and was incorporated into the cells for 2 hours. Total RNA was prepared with (). The EdU-labeled RNAs were then biotinylated and captured by using the Click-it Nascent RNA Capture Kit (Life Technologies), in accordance with the manufacturer’s instructions. Briefly, 1 μg of EU-labeled RNA was biotinylated with 0.5 mM biotin azide in Click-iT reaction buffer. The biotinylated RNAs were precipitated with ethanol at 4°C overnight and resuspended in 50 μl distilled water. The biotinylated RNAs were then mixed with 12 μl Dynabeads MyOne Streptavidin T1 magnetic beads in Click-iT RNA binding buffer and heated at 68°C for 5 min, followed by incubation at room temperature for 30 min while gently vortexing. The beads were immobilized using the DynaMag-2 magnet and were washed with Click-iT wash buffers 1 and 2. The washed beads were resuspended in Click-iT wash buffer 2 and used for cDNA synthesis. cDNA synthesis was performed by the High Capacity cDNA Reverse Transcription Kit as described above.

### Statistical analysis

Experimental replicates of at least 3 for each sample, including negative controls, were prepared whenever applicable. Results are shown as mean ± SD from at least three independent experiments. Statistical analyses were performed using GraphPad Prism 9.4.1 software. Statistical significance was determined by appropriate testing with Student’s t-test and one-way ANOVA with Tukey’s multiple comparison test. A value of p < 0.05 was considered statistically significant.

## Data availability

The RNA-seq, SLAMseq, and CUT and RUN data were deposited in NCBI Gene Expression Omnibus (GEO) database under accession numbers; GSE232843, GSE233119, and GSE233323.

## Acknowledgments

We would like to thank Dr. Robert Yarchoan (NIH/NCI) for the generous gift of rabbit monoclonal antibody against vIL-6. We would like to thank the Flow Cytometry and Genomics Shared Resources at the UC Davis Comprehensive Cancer Center for expert support. This research was supported by public health grants from the National Cancer Institute (CA225266, CA232845) and the National Institute of Allergy and Infectious Disease (AI155515 and AI167663) to Y.I. The Genomics and Flow Cytometry Shared Resources are supported by the UC Davis Comprehensive Cancer Center Support Grant (CCSG) awarded by the National Cancer Institute (NCI P30CA093373).

---

**SFig.1.**
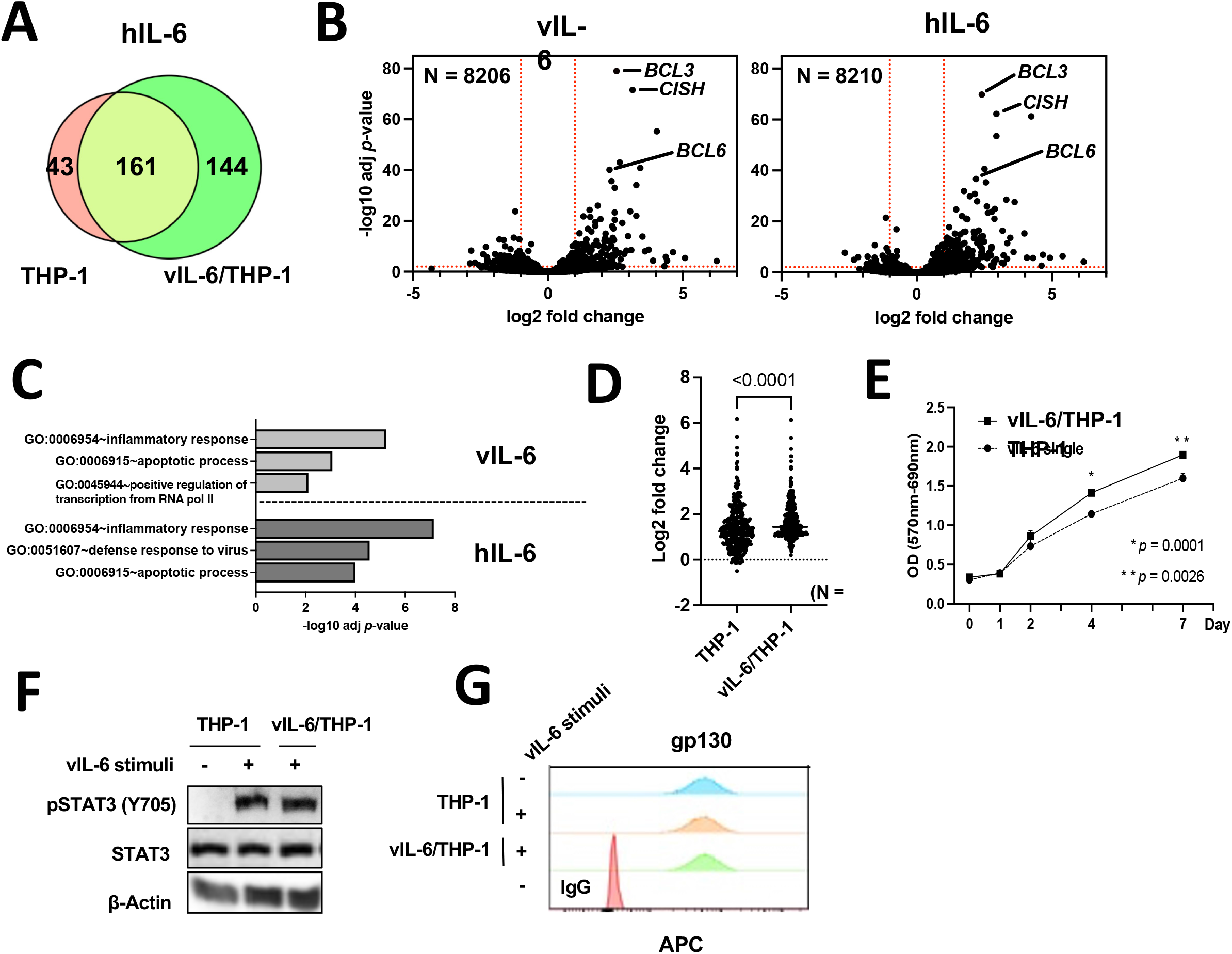
hIL-6 is a functional homolog of vIL-6. **(A)** The number of up-regulated genes (log2 fold change >1, adj *p*-value < 0.01) after hIL-6 stimulation. Red circle represents THP-1 cells and green circle represents vIL-6/THP-1 cells. **(B)** Individual gene expression in THP-1 cells with vIL-6 stimulation (left, N = 8206) and hIL-6 stimulation (right, N = 8210). Representative gene names were labelled adjacent to dots. The red dashed line indicated log2 fold change = ± 1 (vertical) and -log10 adj p-value = 2 (horizontal). **(C)** KEGG pathway analysis performed on up-regulated genes (log2 fold change >1, adj *p*-value < 0.01) in THP-1 cells with vIL-6 and hIL-6 stimulation. The result showed the top three pathways each. **(D)** Individual up-regulated gene expression (N = 348) in parent THP-1 and vIL-6/THP-1 cells after hIL-6 stimulation. Data were analyzed using Wilcoxon matched-pairs signed ranked test and shown as median. **(E)** Measurement of cell proliferation with MTT assays. 1 × 10^4^ THP-1 or vIL-6/THP-1 cells were cultured in triplicate in a 96 well plate. vIL-6 was added to vIL-6/THP-1 cells every other day. OD (570-690nm) was measured on day 0,1,2,4 and 7. Data were analyzed using unpaired Student’s *t* test and shown as mean ± SD. **(F)** Immunoblotting with antibodies directed against STAT3, pshopho-STAT3 (Y705) and β-Actin (loading control) protein in THP-1 and vIL6/THP-1 cells. (G) FACS analysis showing the gp130 expression on cell surface.

**SFig.2.**
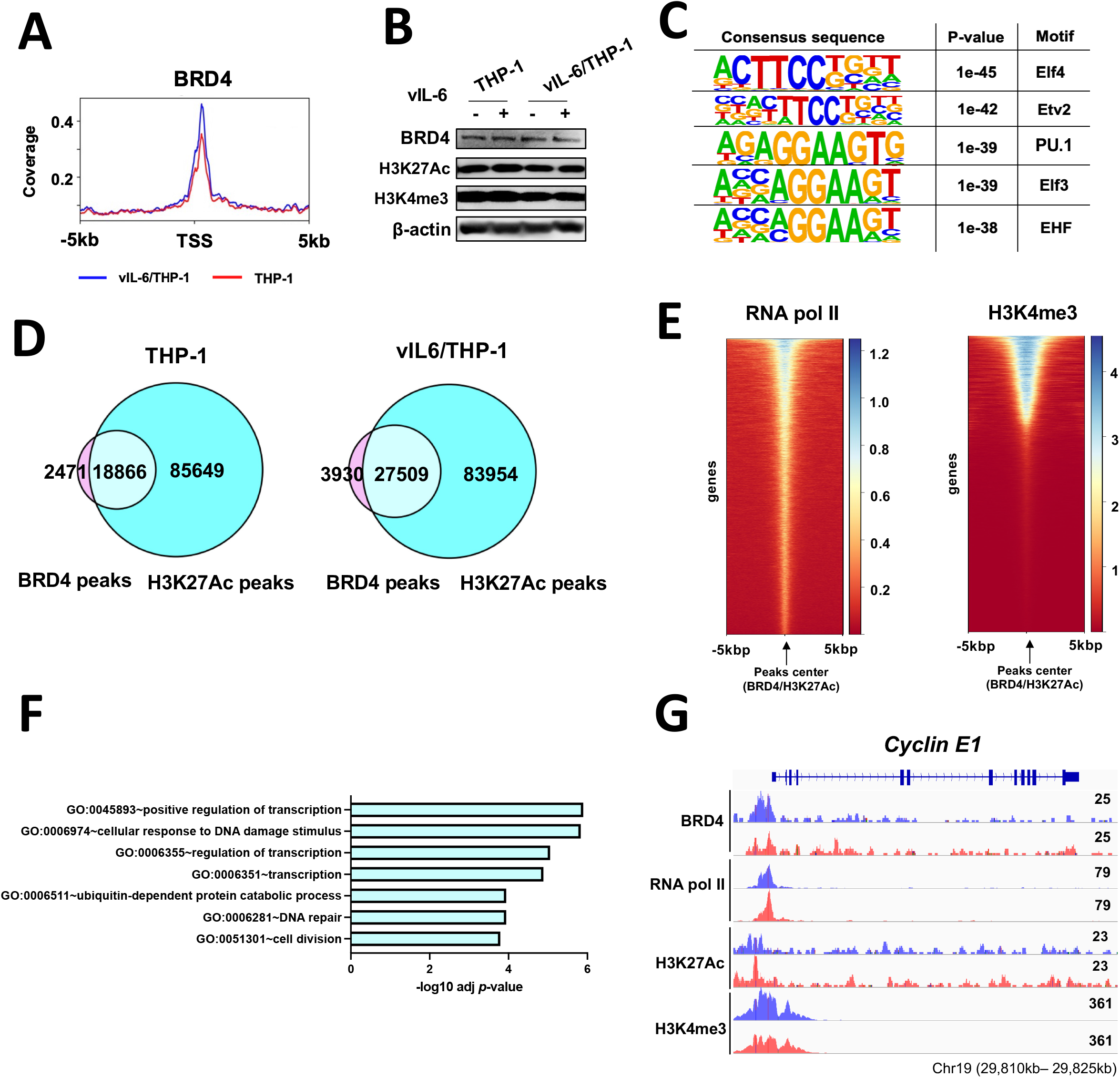
Prolonged vIL-6 exposure enhances the association of BRD4 and H3K27Ac. **(A)** BRD4 CUT &RUN signals in ±5kbp windows around the transcription start sites (TSS) of up-regulated genes in vIL-6/THP-1 cells (N = 303). **(B)** BRD4, H3K27Ac and H3K4me3 protein expression before and after vIL-6 stimulation in parental THP-1 and vIL-6/THP-1 cells. **(C)** DNA binding motif analysis of new BRD4 accumulation sites in vIL-6/THP-1 cells. Images were drawn by findMotif (HOMER). **(D)** The number of BRD4 and H3K27Ac peaks and their association in parental THP-1 and vIL-6/THP-1 cells. The overlapping peaks were extracted using mergepeaks (HOMER). **(E)** RNA pol II and H3K4me3 CUT &RUN signals in ± 5kbp windows around the center of BRD4 and H3K27Ac peaks. **(F)** KEGG pathway analysis performed on genes at BRD4 and H3K27 overlapping peaks in promoter regions in vIL-6/THP-1 cells. Results are presented in descending order. **(G)** BRD4, RNA pol II, H3K27Ac and H3K4me3 enrichment in the *Cyclin E1* promoter region in parental THP-1 cells (pink) and vIL-6/THP-1 cells (blue). The peaks were visualized by importing the BAM files into the Integrative Genomics Viewer (IGV).

**SFig.3.**
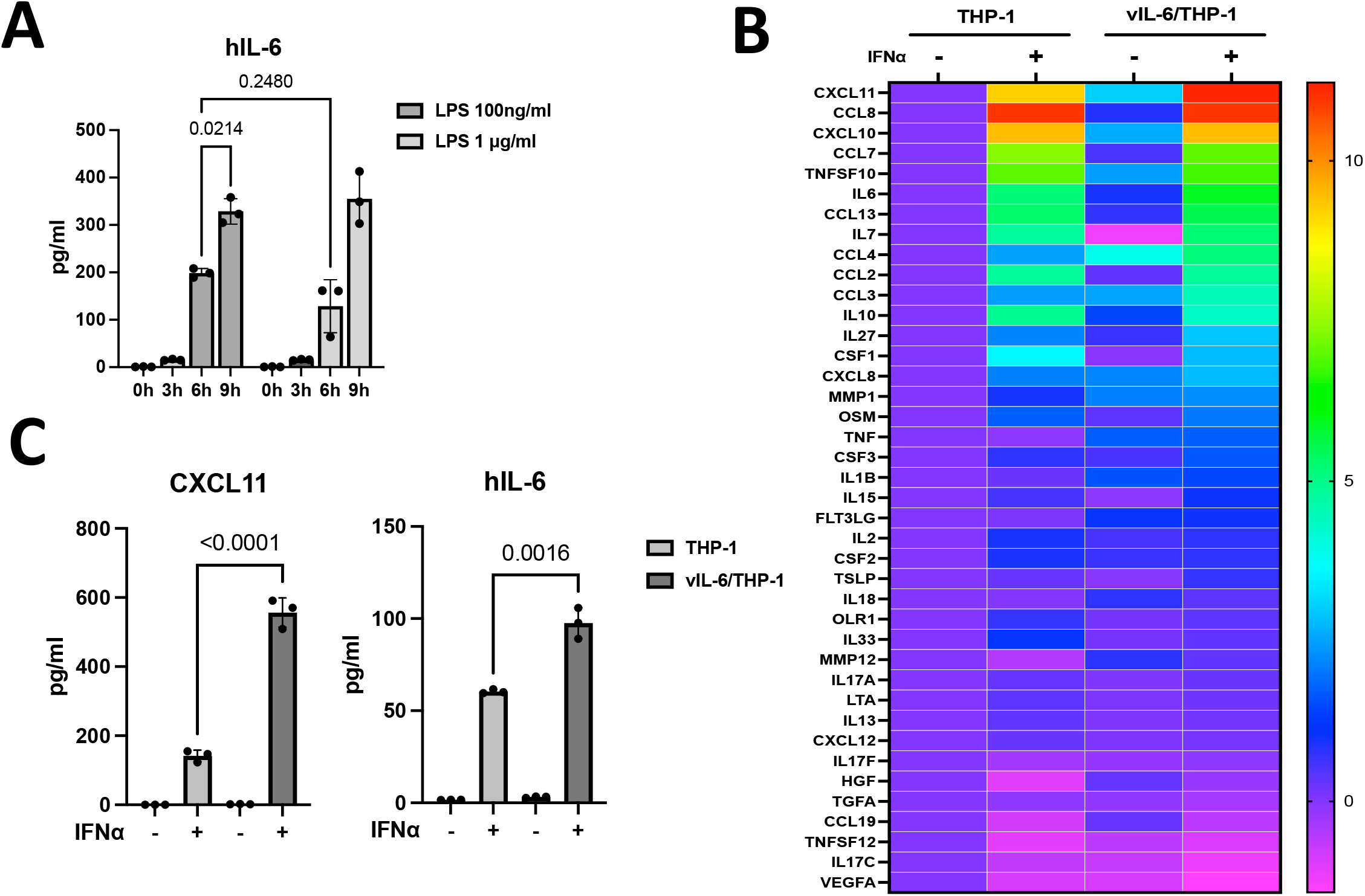
Inflammatory response to IFNα after vIL-6 prolonged exposure. **(A)** hIL-6 production in THP-1 cells by LPS stimulation. THP-1 cells was incubated with LPS 100ng/ml or 1μg/ml for various time periods. Supernatants were harvested and incubated in triplicate in ELISA plate coated with hIL-6 antibody, Human IL-6 Uncoated ELISA kit (Invitrogen) was then used to evaluate the hIL-6 production by following the manufacturer’s guideline. The protein binding measured as OD values at 450nm was shown. Results are presented as mean percentage viability ±SD *(n = 3* samples/group). Data was analyzed by a one-way ANOVA test. **(B)** Heatmap showing the results of Olink® Target 48Cytokine panel . IFNα (100 n g/ m l) was added toparent THP-1 cells or vil-6/THP-1cells for6 hours. Cytokine production in untreated THP-1 cells was set as 1 andlog_2_ foldactivation relative to untreated cells are shown. Samples were prepared in triplicate and the mean value were shown. **(C)** Inflammatory cytokine production determined by Olink proximity extension assay. Data was analyzed using two-sided unpaired Student’s *t* testandshownas mean ± SD.

**SFig.4.**
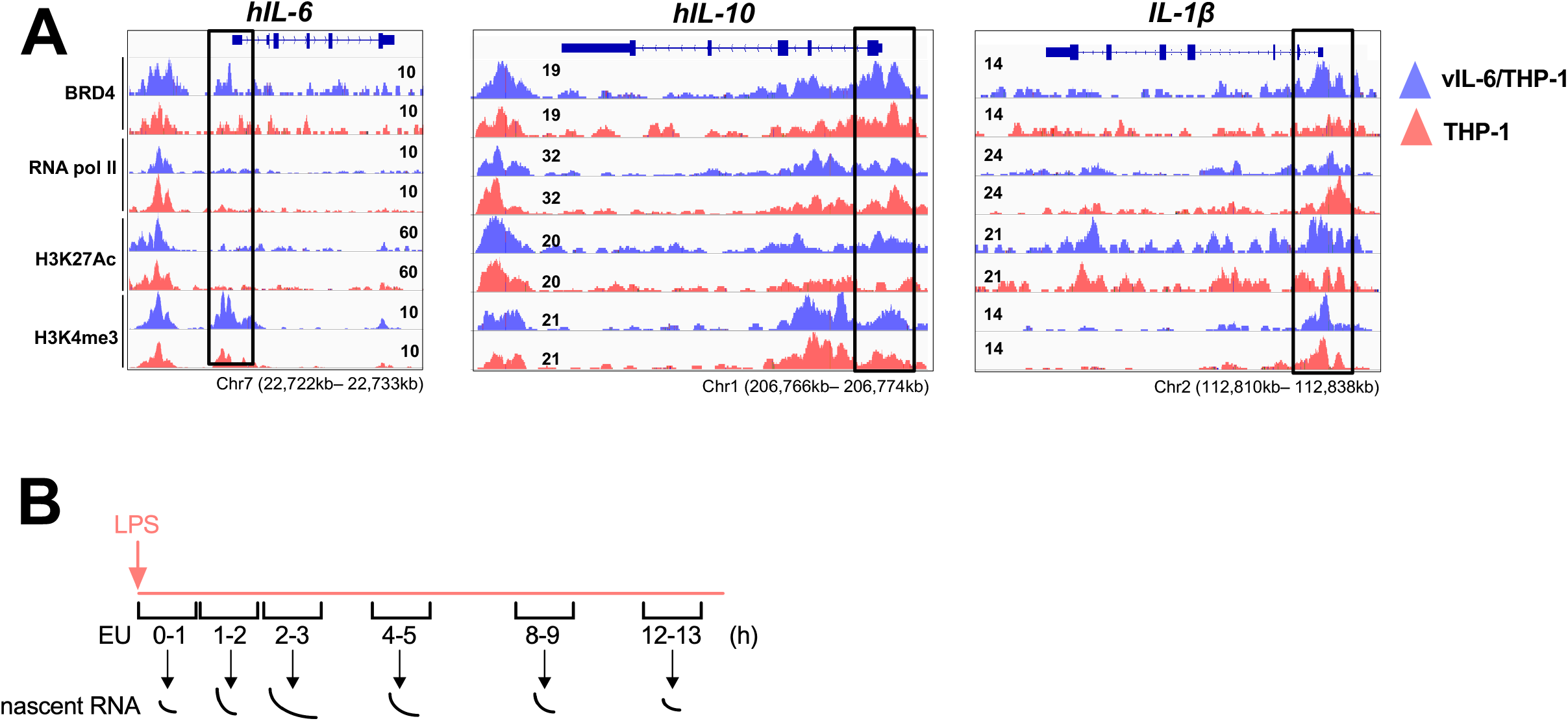
BRD4 enrichment in the promoter region of inflammatory genes. **(A)** BRD4, RNA pol II, H3K27Ac and H3K4me3 enrichment in hIL-6, IL-10, IL-1β promoter region in parental THP-1 cells (pink) and vIL-6/THP-1 cells (blue). The promoter region is enclosed by a black line. Each CUT&RUN peak was visualized by importing the BAM files into Integrative Genomics Viewer (IGV). **(B)** Schematic diagram of nascent RNA labeling after LPS stimulation. LPS were added to culture medium in parental THP-1 and vIL-6/THP-1 cells and cells were incubated with EU for1 hour in0,1,2,48, 12 h LPS stimulation.

**Table. S1.**
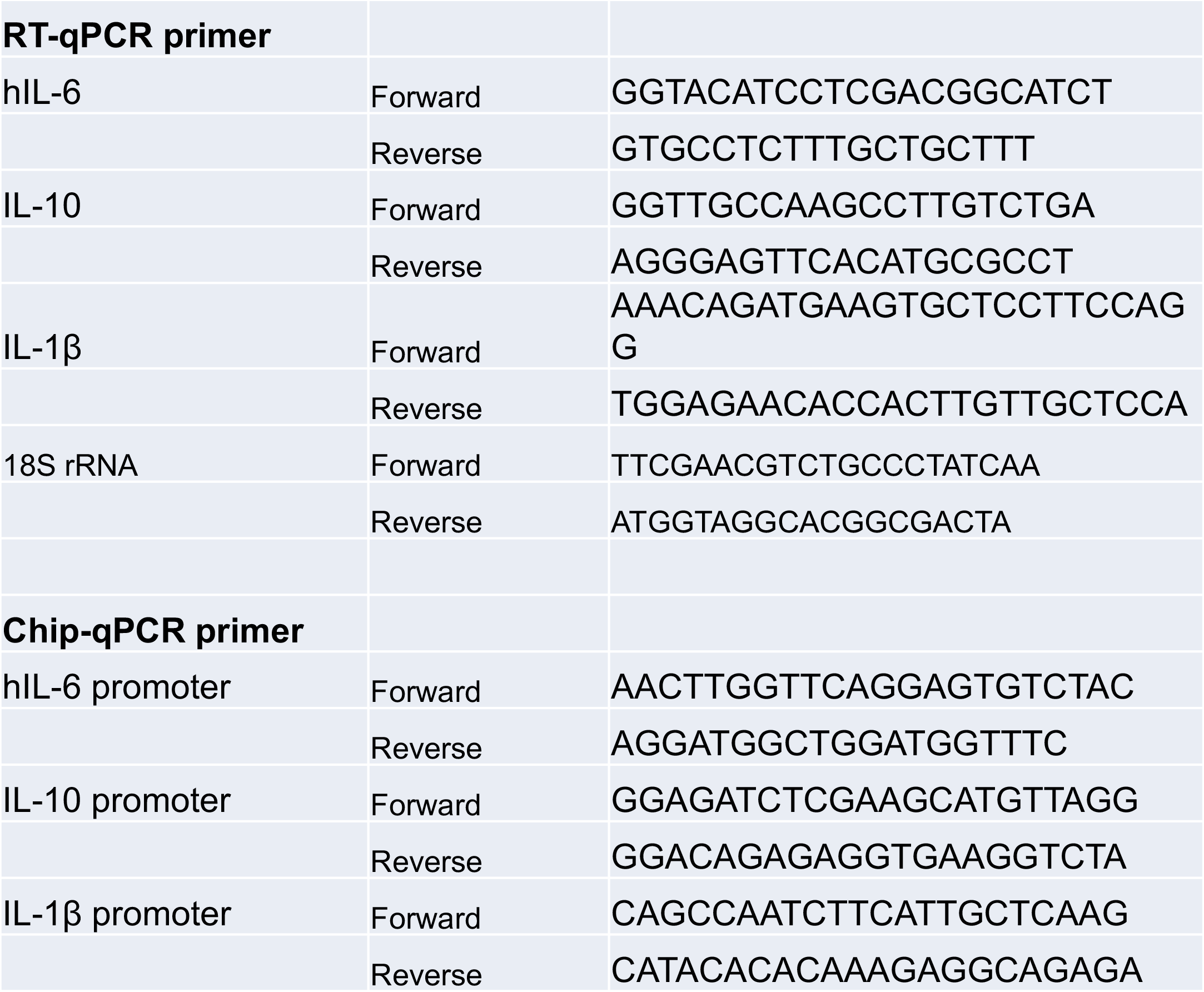
Primer used for RT-qPCR and Chip-qPCR

